# Multi-parameter phenotyping of platelets and characterisation of the effects of agonists using machine learning

**DOI:** 10.1101/2023.12.08.570628

**Authors:** Ami Vadgama, James Boot, Harriet E. Allan, Charles A. Mein, Paul C. Armstrong, Timothy D. Warner

**Author notes:** PCA and TDW contributed equally to this work. Corresponding author: Dr Paul C. Armstrong, Centre for Immunobiology, Blizard Institute, Barts and the London School of Medicine and Dentistry, Queen Mary University London, 4 Newark Street, London, E1 2AT, United Kingdom.

## Abstract

**Background:** Platelets are crucial for thrombosis and haemostasis, with their function driven by the expression of specialised surface markers. The concept of distinct circulating sub-populations of platelets has emerged in recent years, but their exact nature remains debatable. We reasoned that a more comprehensive characterisation of surface marker changes at rest and upon activation would be valuable in determining this.

**Objective:** To use a full spectrum flow cytometry-based panel, together with parameters of physical properties, to describe surface marker changes in healthy platelets at rest and on activation, and to observe how these responses differ according to platelet age.

**Methods:** A 14-marker flow cytometry panel was developed and applied to vehicle- or agonist-stimulated platelet-rich plasma samples obtained from healthy volunteers, or to platelets sorted according to SYTO-13 staining intensity as an indicator of platelet age. Data were analysed using both user-led and independent approaches incorporating novel machine learning-based algorithms.

**Results:** The assay detected changes in marker expression in healthy platelets, at rest and on agonist activation, that are consistent with the literature. Machine learning identified stimulated populations of platelets with high accuracy (>80%). Similarly, differentiation between young and old platelet populations achieved 76% accuracy, primarily weighted by FSC-A, CD41, SSC-A, GPVI, CD61, and CD42b expression patterns.

**Conclusions:** Our findings provide a novel assay to phenotype platelets coupled with a robust bioinformatics and machine learning workflow for deep analysis of the data. This could be valuable in characterising platelets in disease. *(240 words)*

**Essentials:**

- Platelet function is directed by the expression of specialised surface markers
- Circulating platelet sub-populations are incompletely characterised
- Multi-parameter spectral flow cytometry allows robust and comprehensive phenotyping of platelets
- Coupling multi-parameter spectral flow cytometry with machine learning offers a powerful method to determine platelet sub-populations

## Introduction

Haemostasis is a carefully orchestrated process in which platelets are primary players. These metabolically-active cell fragments circulate for approximately 7-10 days in healthy individuals before being degraded by the spleen or liver[1–3]. Platelet function is mediated through the expression of specialised surface markers, with resting and activated platelets showing different expression profiles consistent with response heterogeneity in thrombus formation[4,5]. These variations in surface marker expression are thought to be conferred during production from megakaryocytes, activation history, and ageing in the circulation[6,7], and may describe dynamically discrete and specialised platelet sub-populations[8,9].

As platelets age in the circulation, they lose messenger ribonucleic acids (mRNA) remaining from their progenitor megakaryocytes[10–12]. Platelets have minimal capacity to make new mRNA, and thus these residual mRNAs can be used as a surrogate measure for age[13–15]. Newly-formed or ‘young’ platelets (also termed reticulated platelets or the immature platelet fraction) have the highest levels of mRNA, while ‘old’ platelets have the lowest[14,16,17]. Young platelets are hyper-reactive and have an elevated thrombotic potential[18–20]. This is apparent in several pathological states, such as diabetes mellitus[21] and major trauma[18], in which there are relative increases in young platelets resulting from altered platelet turnover and lifespan associated with increased incidences of thromboembolic events[22–25] and decreased efficacy of anti-platelet therapies[26–29].

Flow cytometry with the inclusion of fluorescently-tagged antibodies is frequently used to investigate platelet protein expression and function as it requires only small volumes of blood and a relatively low number of platelets, making it ideal for analysis of clinical samples. However, the number of parameters that can be measured concurrently using this method is limited by overlap of fluorescent emission spectra, resulting in antibody panels that typically determine, at most, three to four markers in any one sample [30,31]. Spectral flow cytometry is a next-generation technique that allows the simultaneous measurement and discrimination of multiple fluorophores by evaluation of full emission spectral signatures. Accounting for steric hindrance, this platform can therefore be used for analysis of 10-15 markers on platelets. Previous work from Blair *et al*.[8] has used mass cytometry to develop a panel to study platelet function, but spectral flow cytometry offers several considerable advantages including cost of reagents, availability of equipment, and simplicity of technique. With this in mind, we developed a 14-marker spectral flow cytometry panel based upon the panel of Blair *et al*. (noting in this report that Blair *et al*. have subsequently published reports utilizing spectral flow cytometry[30,32]). In our studies we subjected our data to a powerful computational analytical approach employing machine learning to explore the potential existence of platelet sub-populations in the human circulation with a particular focus on young and old platelets.

## Materials and Methods

### Ethical statement: human studies

All studies were conducted according to the principles of the Declaration of Helsinki and were approved by St. Thomas’ Hospital Research Ethics Committee (Ref.: 07/Q0702/24). Healthy volunteers were aged 18-40, screened prior to entering the study (non-smokers; had not taken non-steroidal anti-inflammatory drugs ≤10 days prior to donating blood; no health problems contraindicating study involvement) and gave written informed consent.

### Collection of blood and preparation of platelet-rich plasma

Blood was drawn from volunteers by venepuncture into trisodium citrate vacutainers (3.2%; BD Biosciences), and platelet-rich plasma (PRP) was isolated as previously published[24].

### Flow cytometric measurement of activation markers

PRP was diluted 1:40 with 2mM Ca^2+^-buffered, filtered PBS, and added to a 96-well plate with wells containing vehicle (phosphate-buffered saline, PBS) or agonist: 0.3-30μM adenosine diphosphate (ADP; Labmedics); 0.3-30μM thrombin-receptor activating peptide 6 (TRAP-6; Cambridge Biosciences); 0.3-30μM U46619 (Enzo Life Sciences); 3-100μM protease-activated receptor 4 agonist (PAR-4; Cambridge Biosciences); 0.03-3μM collagen-related peptide (CRP-XL; University of Cambridge). In initial experiments, an antibody mix comprising anti-CD42b-BV421 (1:70; clone HIP1; BioLegend), PAC-1-FITC (1:10; BD Biosciences), and anti-CD62P-APC (1:100; clone AK4; BioLegend) was added to each well. In later experiments, an antibody master mix (Supplementary table 1) and staining buffer (Brilliant Stain Buffer, BD Biosciences) were used. The plate was then mixed (200rpm; 37°C) for 20 minutes in the dark (BioShake iQ, Quantifoil Instruments GmbH). Samples were fixed with 1% formalin and run on the Cytek Aurora 5-laser flow cytometer (Cytek Biosciences). Platelets were gated on SSC-A/CD42b-A, and 10,000 CD42b+ events were collected.

### Flow cytometric sorting of young and old platelets

Adapting our previously published approach[33], PRP was stained with SYTO-13 (750nM; Thermo-Fisher Scientific) prior to sorting. 10 million platelets per condition were sorted using a BD FACS Aria IIIu Fusion Cell Sorter (70μM nozzle, 70Ps, ≤10,000 events/second; BD Biosciences) with the top 20% SYTO-13 fluorescence being taken as ‘young’ and the bottom 30% SYTO-13 as ‘old’. Platelets were pelleted in the presence of prostacyclin (Epoprostanol; 2μmol/L; Tocris Biosciences) at x1000*g* for 10 minutes and re-suspended in calcium chloride (CaCl_2_; 2mmol/L; Sigma-Aldrich) -buffered MTH buffer. Panel markers were then measured as described above using the Cytek Aurora.

### Statistical, bioinformatics, and machine learning analysis

Data were collected using SpectroFlo v2 (Cytek Biosciences) software, analysed using NoVo Express 1.3.0 (ACEA Biosciences Inc.), FlowJo v10 (TreeStar Inc.), and GraphPad Prism 9 (GraphPad Software Inc.) software. Data are expressed as median fluorescence intensity (MFI) ±SEM, and analysed with a one-/two-way ANOVA or mixed-effects analysis followed by multiple comparison post-hoc tests, as appropriate. Correlations were assessed by non-linear regression. Statistical significance was assumed for p <0.05.

Bioinformatic analysis was performed using R version 4.2 or later. Data were loaded into an RStudio (2022.02.2) environment using a loading function from the flowCore (v2.10.0 or later) package[34,35]. Data loading checks for quality control purposes were performed by checking the correlations between MFIs of all parameters in the loaded data and FlowJo v10-analysed data. Data were further analysed using the Spectre package[36] and associated functions. Namely, the data was transformed using the logicle transformation and then dimensionality reduction was performed using principal component analysis (PCA) followed by t-distributed stochastic neighbour embedding (tSNE: an unsupervised, non-linear dimensionality reduction technique for visualising high-dimensional data in a two- or three-dimensional space). Individual platelets are represented as single points and grouped together based on their degree of similarity of expression patterns of all 14-16 parameters. tSNE also identifies heterogeneity in platelet responses, allowing the identification of sub-populations of platelets. FlowSOM, a self-organising map clustering algorithm, was used for cluster generation, which automatically determines the optimal number of clusters for the dataset.

Using the Caret v6.0-93 (Classification And REgression Training) R package[37], we developed a machine learning model to predict whether platelets were treated with vehicle or agonist. Platelet data were loaded and processed as already detailed, with 10,000 platelets per healthy donor per treatment being loaded, unless otherwise stated. Balanced in number for each condition, from 16 donors, a Random Forest model was given a training dataset made up of 80% of the total data to learn from. The remaining 20% of the total data was used for preliminary validation. The remaining 5 donors were used as an “unseen” validation data set to test the model. A feature importance comparison was then run to determine which markers were most important in the classification. The model was trained using a 10-fold cross-validation using 3 repeats. This means that the training data is first randomly shuffled and split into 10 “folds”, then, in turn, each fold is excluded from the training data whilst the remaining 9 folds are used to train a model; the excluded fold is used to assess the accuracy of the trained model. This process was then repeated a total of 3 times; each time the data was shuffled randomly, producing new folds. The metric used to assess the quality of the model was the Receiver Operating Characteristic (ROC); overall accuracy was also reported.

Using the Caret v6.0-93 (Classification And REgression Training) R package[37] we developed Random Forest machine learning models to predict whether platelets were young or old in the presence of vehicle or individual agonists. Platelet data were loaded and processed as already detailed, with 9,000 platelets per young and per old sample from each donor being loaded, outlier events/platelets were removed, and the measurements for PAC-1 excluded. In total there were 8 donors, each with a young and old sample. 6 of the 8 donors were used to create a training and preliminary validation dataset; 80% of the data from the 6 donors was used for training whilst 20% was used for preliminary validation to assess the accuracy of the model. Data from the remaining 2 donors was kept aside to use as an “unseen” validation dataset. As previously, a feature importance comparison was then run to determine which markers were most important in the classification. The model was trained using a 10-fold cross-validation using 3 repeats, and ROC used to assess the quality of the model, as already detailed.

## Results

### Assay capable of detecting significant changes in marker expression on activation in healthy platelets

To select optimum concentrations of agonists for use in subsequent experiments, flow cytometry was used to assess changes in PAC-1 binding and CD62P expression in response to increasing concentrations of TRAP-6, PAR-4, CRP-XL, ADP, and U46619 (Supplementary figure 1). The following concentrations were selected for their ability to induce a robust response: 10µM TRAP-6, 100µM PAR-4, 3µM CRP-XL, 30µM ADP, and 10µM U46619.

The chosen concentrations of agonists were then tested using the extended phenotyping panel of markers as well as forward scatter (FSC-A) and side scatter (SSC-A). All agonists tested caused significant increases in the expression of CD62P, PAC-1, CD63, CD107a, CD61, CD29, and CD9, and decreases in CD42b; none of the agonists caused any changes in CLEC-2 (Figure 1). TRAP-6 was the only agonist to decrease CD31 (4336±233 vs. 3973±223, p=0.03), and CRP-XL was the only agonist to decrease GPVI (5735±512 vs. 4030±600, p=0.006). CD42a expression was decreased by all agonists except CRP-XL.

**Figure 1:**
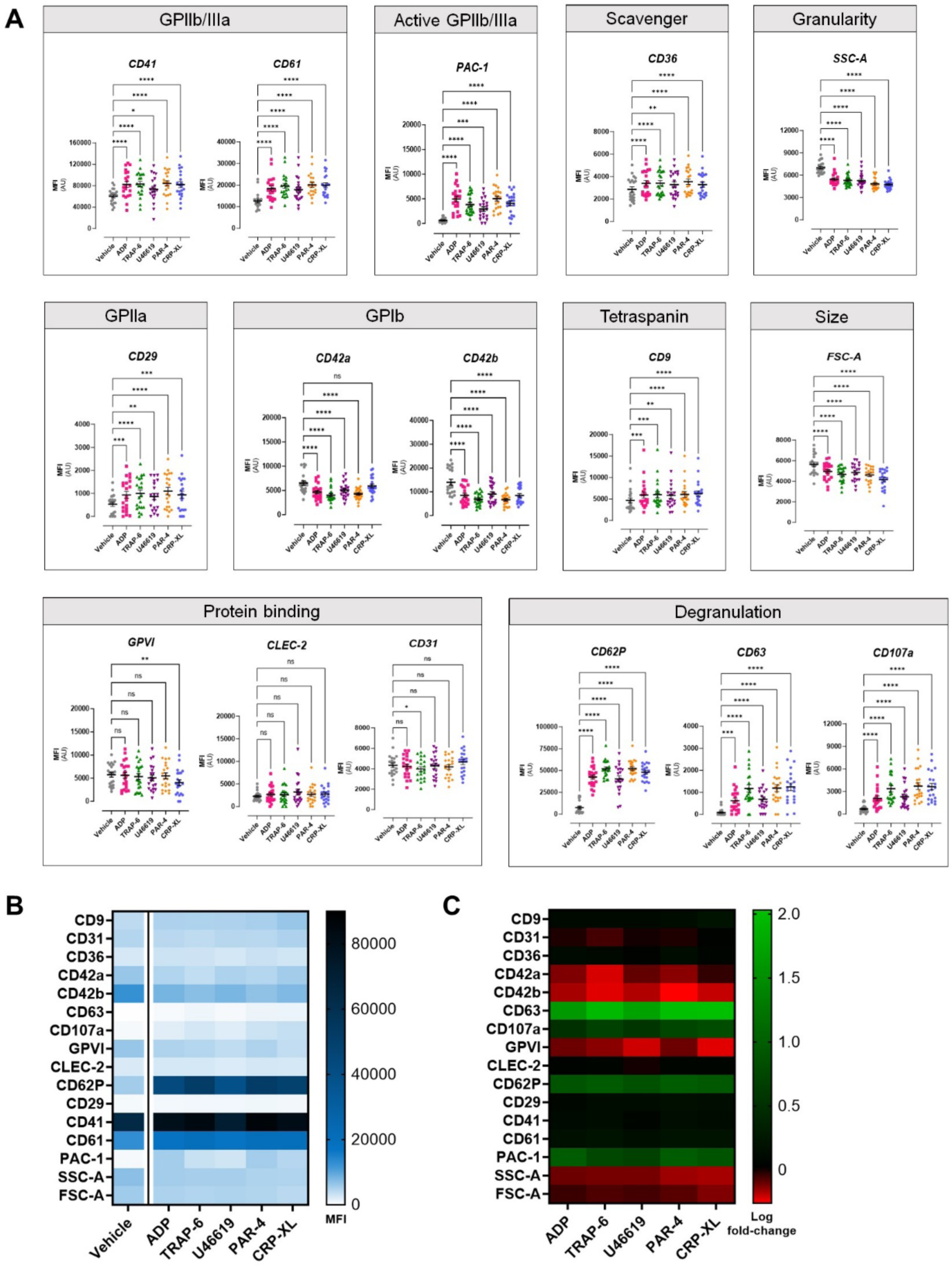
Changes in individual surface marker expression in resting and activated platelets expressed as (A) raw MFI ±SEM, (B) a heatmap based on MFI values, and (C) a heatmap based on MFI log fold-changes. Data were analysed using a one-way ANOVA with a Dunnett test to correct for multiple comparisons (n=20-21).

### High-dimensionality analysis allows visualisation and in-depth interrogation of marker changes in response to activation

Unsupervised dimensionality reduction and visualisation of the entire data sets using tSNE revealed the same shifts in receptor patterns. Namely, agonist stimulation caused visible increases in the expression of PAC-1, CD62P, CD63, CD107a, CD61, and CD9, decreases in CD42b, CD42a, GPVI, CLEC-2, and CD31; CD29 remained unchanged (Figure 2). The same visualisation approach also confirmed that the detected marker expression patterns were shared across donors and not driven by donor or batch effect (Supplementary figure 2). Auto-clustering analysis produced 5 clusters in vehicle-treated platelets and between 9 and 11 clusters in agonist-treated platelets. Each formed cluster was not dominated by individual donors but rather reflected the gradation in difference of expression (Supplementary figure 3). However, hierarchical dendrograms within the clustering indicated that interlinked relationships between the expression of each marker differed by agonist stimulation (Supplementary figure 3).

**Figure 2:**
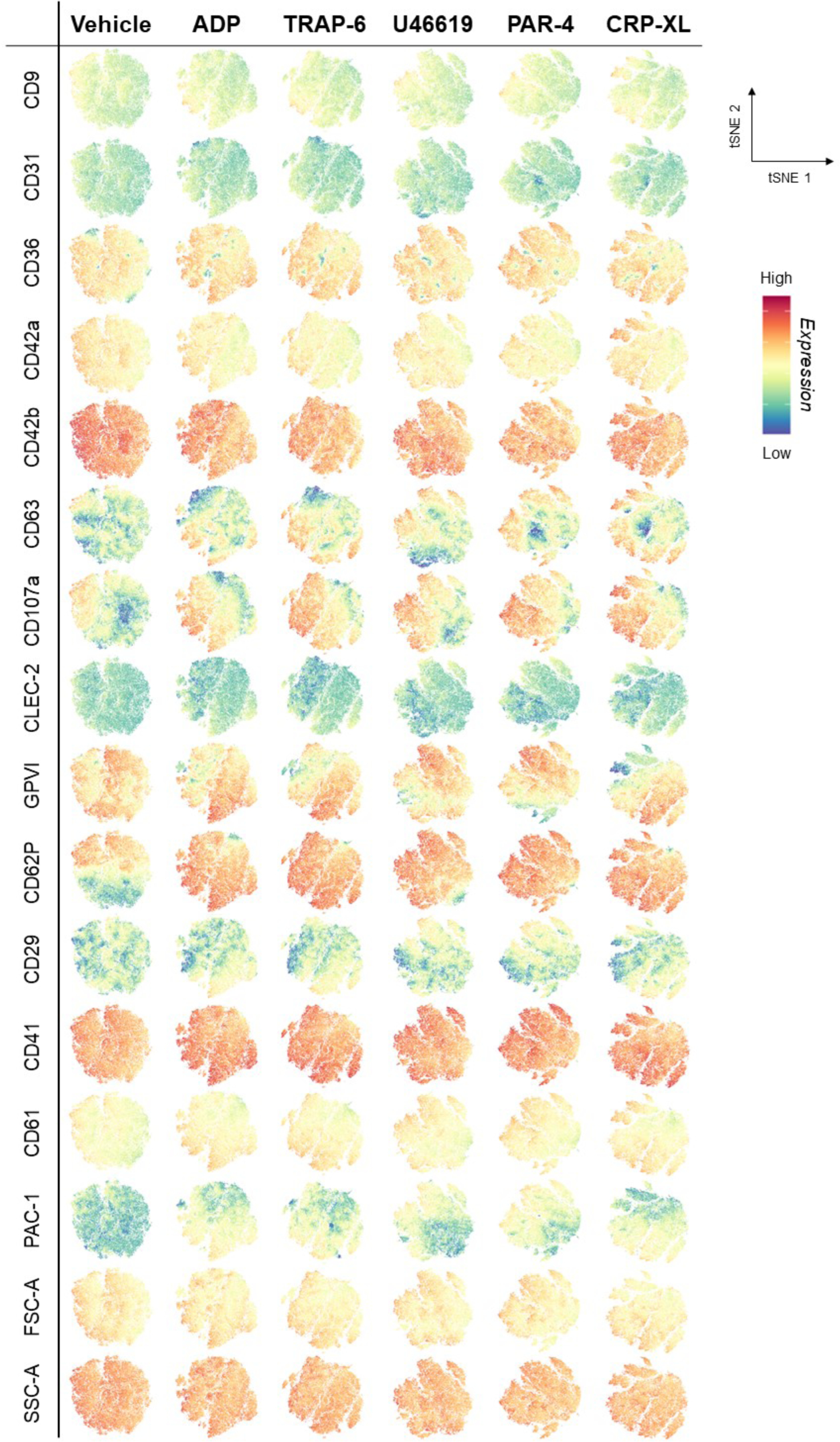
High dimensionality analysis of platelet sub-populations at rest and in agonist-activated platelets. Colour intensity correlates with marker expression (low=blue, high=red). Data were analysed using tSNE (n=20-21).

### Machine learning reveals most the important markers in distinguishing effects of agonists

Machine learning (ML) was used to further analyse the data at the single platelet level in an unbiased fashion. Following training, the accuracy of differentiation of vehicle-treated from agonist-treated platelets was determined. Accuracy rates for unseen datasets were highest for TRAP-6, PAR-4, and CRP of 0.92, 0.91, and 0.88, respectively. Comparatively, rates for ADP and U46619 were 0.80 and 0.77, respectively (Table 1), with the greatest fall between the training and unseen sets also occurring with these agonists.

**Table 1:** Predictive efficacy of machine learning of stimulated platelets.

|  | **ADP** | **TRAP-6** | **U46619** | **PAR-4** | **CRP-XL** |
| --- | --- | --- | --- | --- | --- |
| Training set accuracy | 0.92 | 0.94 | 0.89 | 0.95 | 0.95 |
| Unseen set accuracy | 0.80 | 0.90 | 0.78 | 0.87 | 0.89 |

Rankings of distinguishing markers within each prediction model was examined with those with a weighting of greater than 20 considered important (Figure 3). For identification of TRAP-6-, PAR-4-, and CRP-XL-stimulated platelets, CD62P was considered the most important, followed by PAC-1, CD42b, and CD107a (Figure 3A). Identification of ADP-treated platelets was based predominantly on the expression patterns of PAC-1, CD62P, and CD42b (Figure 3B-D). This was similar to platelets activated with U46619, for which CD62P, CD42b, and PAC-1 were highest (Figure 3E).

**Figure 3:**
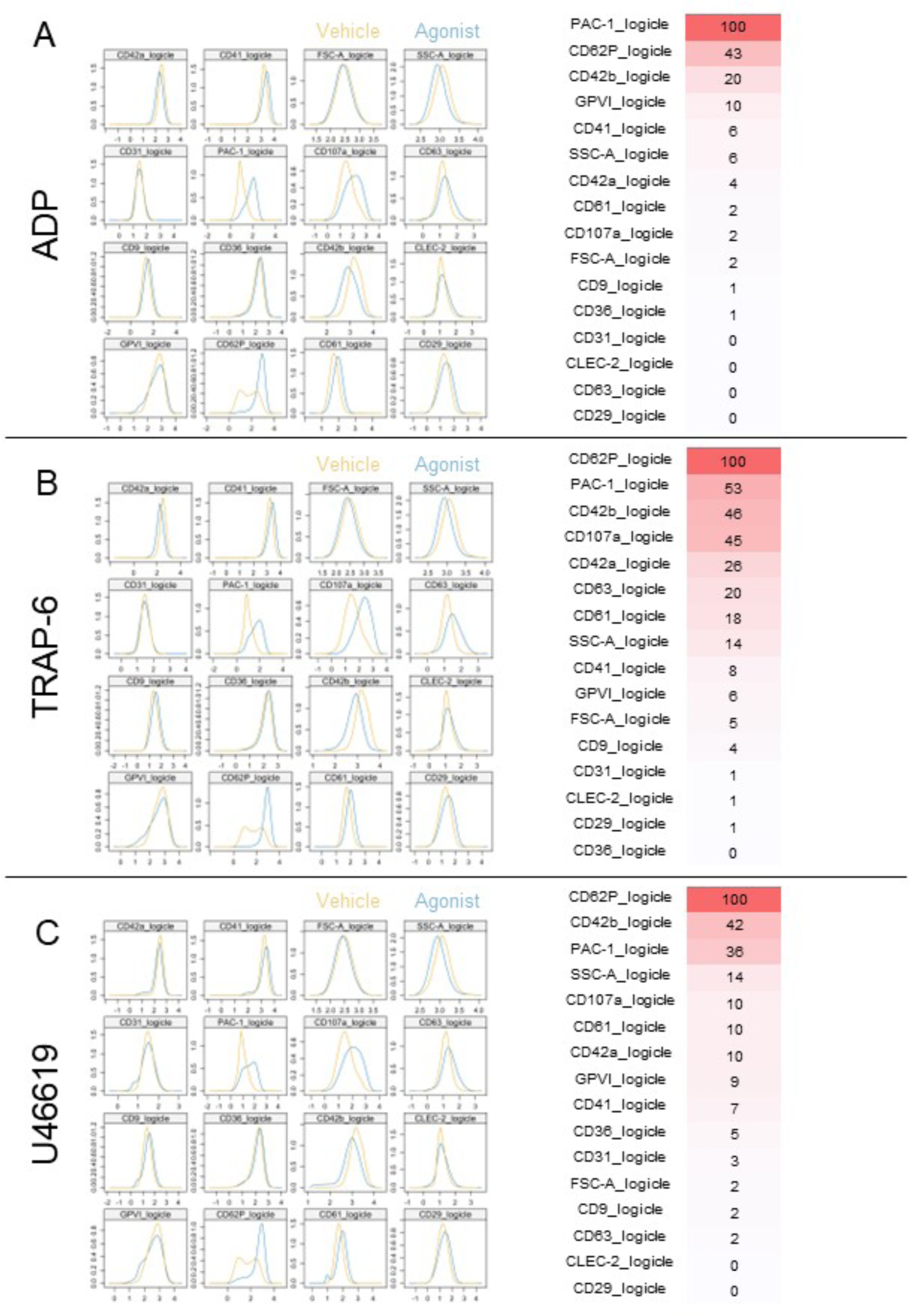
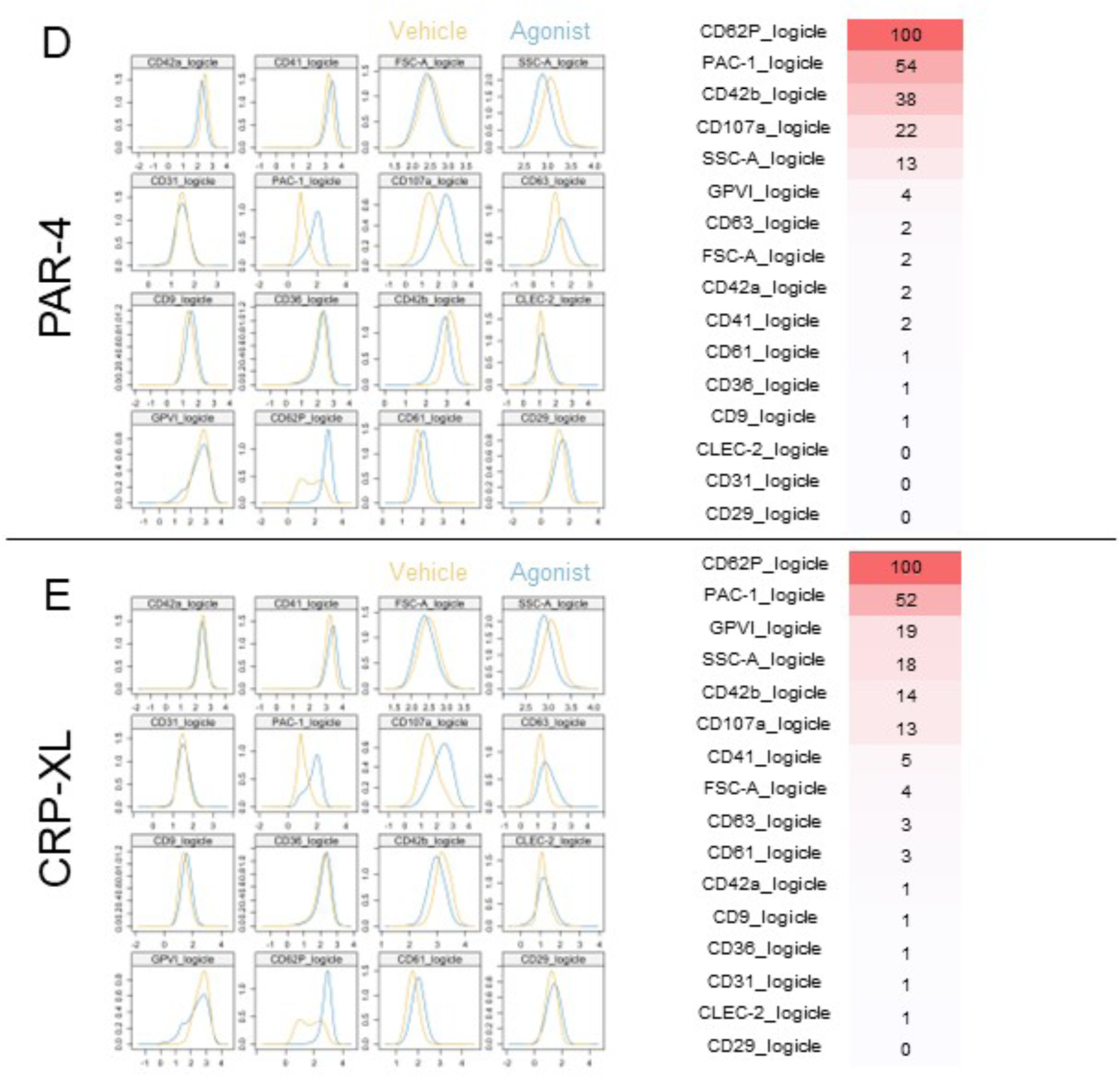
Markers used by machine learning algorithm to distinguish between vehicle- and (A) ADP-treated, (B) TRAP-6-treated, (C) U46619-treated, (D) PAR-4-treated, and (E) CRP-XL-treated platelets, listed in order of importance (most important=red, least important=white; n=20-21).

### CD41/CD61, FSC-A, SSC-A, GPVI, CLEC-2, and CD61 are the primary markers used by machine learning to differentiate between young and old platelets

Next, we undertook phenotypic analysis of young and old platelets, as determined by SYTO-13 RNA staining, following vehicle or agonist stimulation. We first analysed the data on a traditional by-population basis by looking at raw MFI values and changes in MFI (Supplementary figure 4), before applying machine learning. Predictive capability within each training set ranged from 0.86 to 0.90, however decreased in the unseen datasets to range from 0.74 to 0.78 (Table 2).

**Table 2:** Predictive efficacy of machine learning of young and older platelets.

|  | **Vehicle** | **TRAP-6** | **PAR-4** | **CRP-XL** | **ADP** | **U46619** |
| --- | --- | --- | --- | --- | --- | --- |
| Training set accuracy | 0.86 | 0.87 | 0.86 | 0.86 | 0.90 | 0.86 |
| Unseen set accuracy | 0.76 | 0.78 | 0.77 | 0.76 | 0.76 | 0.74 |

Markers rated greater than an importance of 20 within vehicle-treated platelets identified FSC-A, CD41, SSC-A, GPVI, and CD61 as most important (Figure 4). FSC-A, SSC-A, and CD41 were the top three discriminators for all agonists tested, with CD61, CD42b, and GPVI being also seen at above 20 in all. CLEC-2 was seen in TRAP-6, and CD61 in CRP-XL.

**Figure 4:**
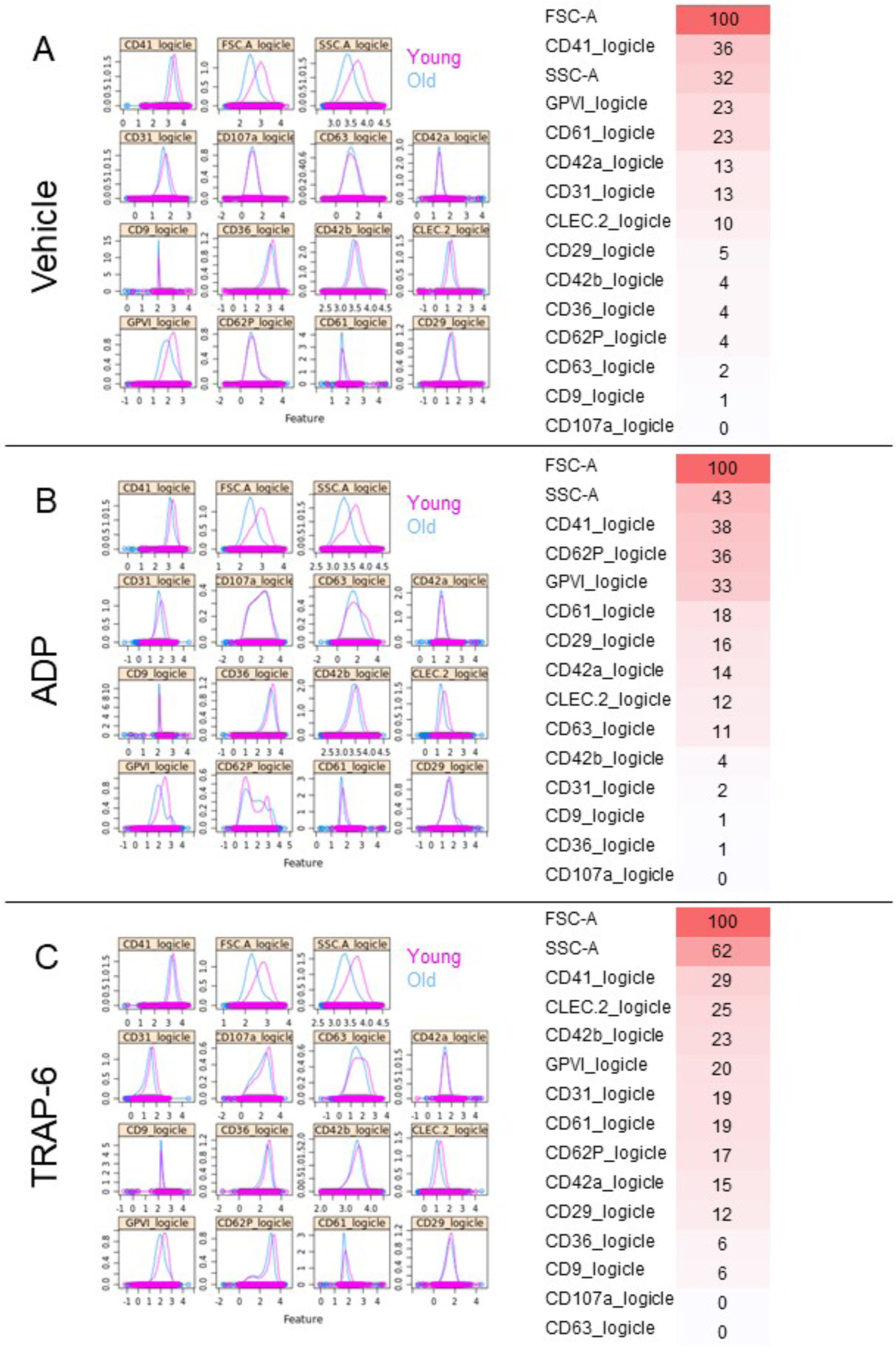
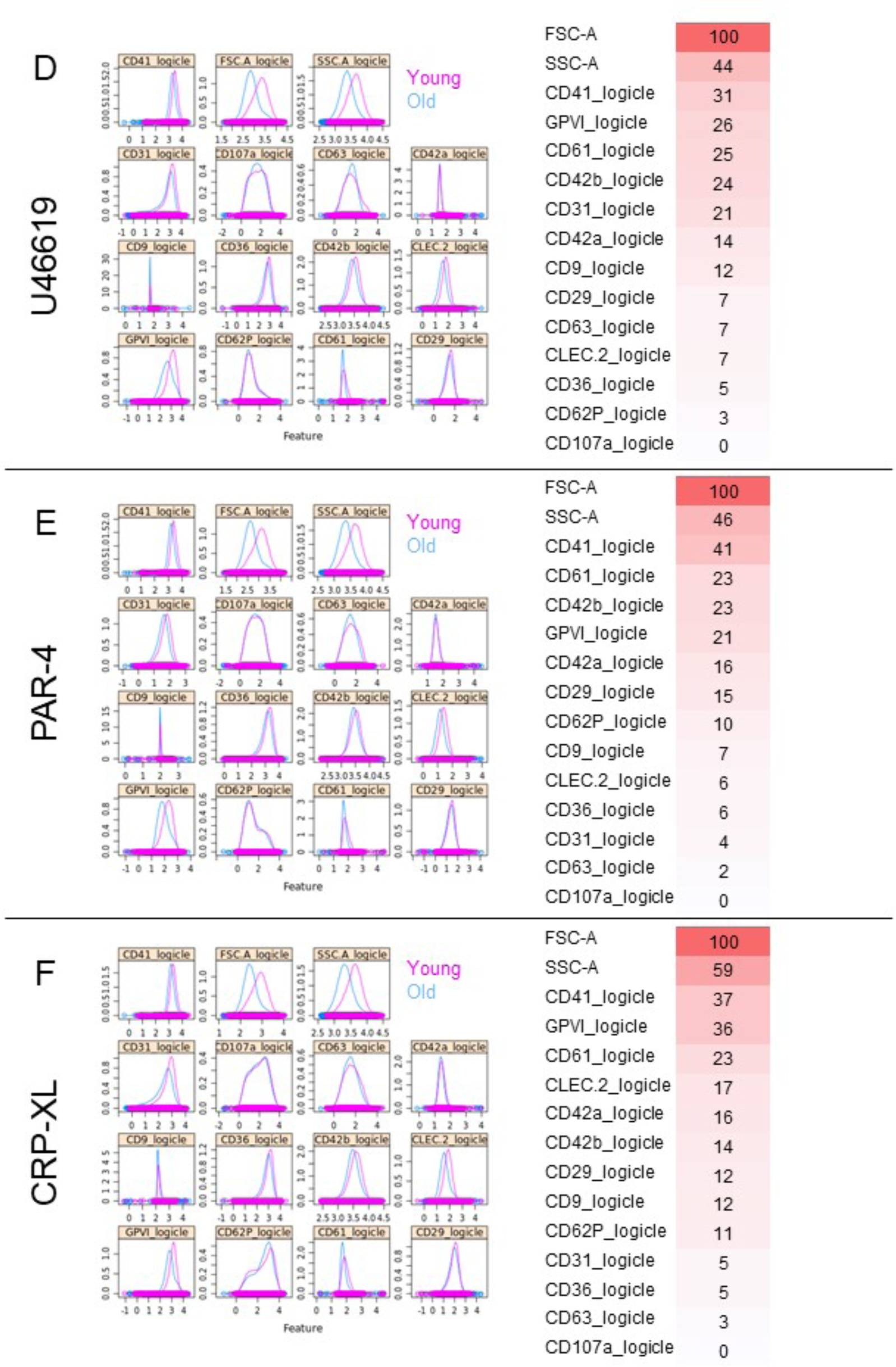
Markers used by machine learning algorithm to differentiate between (A) vehicle-, (B) ADP-, (C) TRAP-6-, (D) U46619-, (E) PAR-4-, and (F) CRP-XL-treated “young” and “old” platelets, listed in order of importance (most important=red, least important=white; n=8).

Finally, we applied the markers rated greater than an importance of 20 from our separated young and old platelet populations to the mixed healthy platelet data presented previously. This analysis indicated an increased probability of young platelets being present in clusters with higher markers of activation, both in control conditions (i.e. vehicle-treated) and following exposure to platelet agonists (Figure 5).

**Figure 5:**
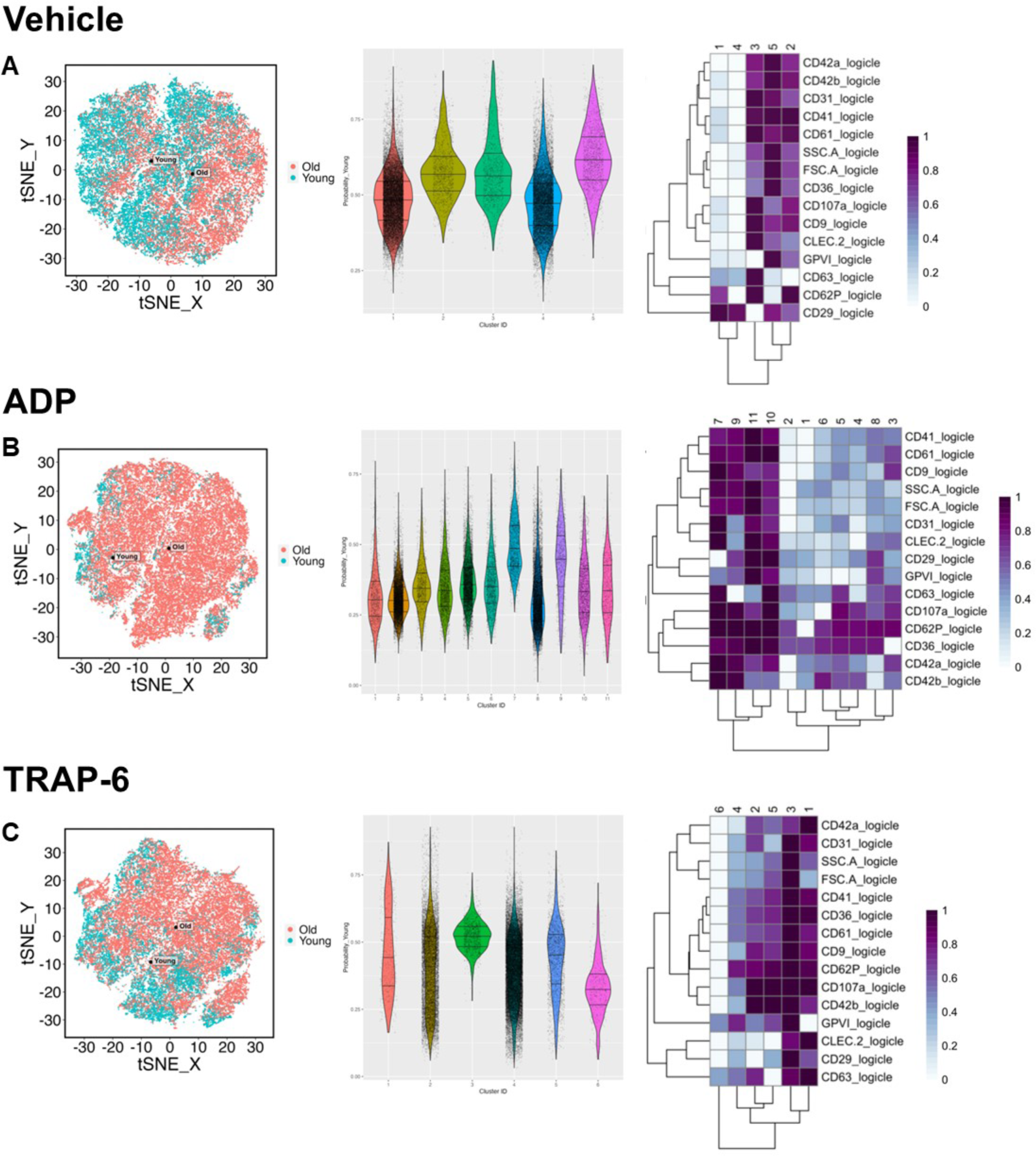
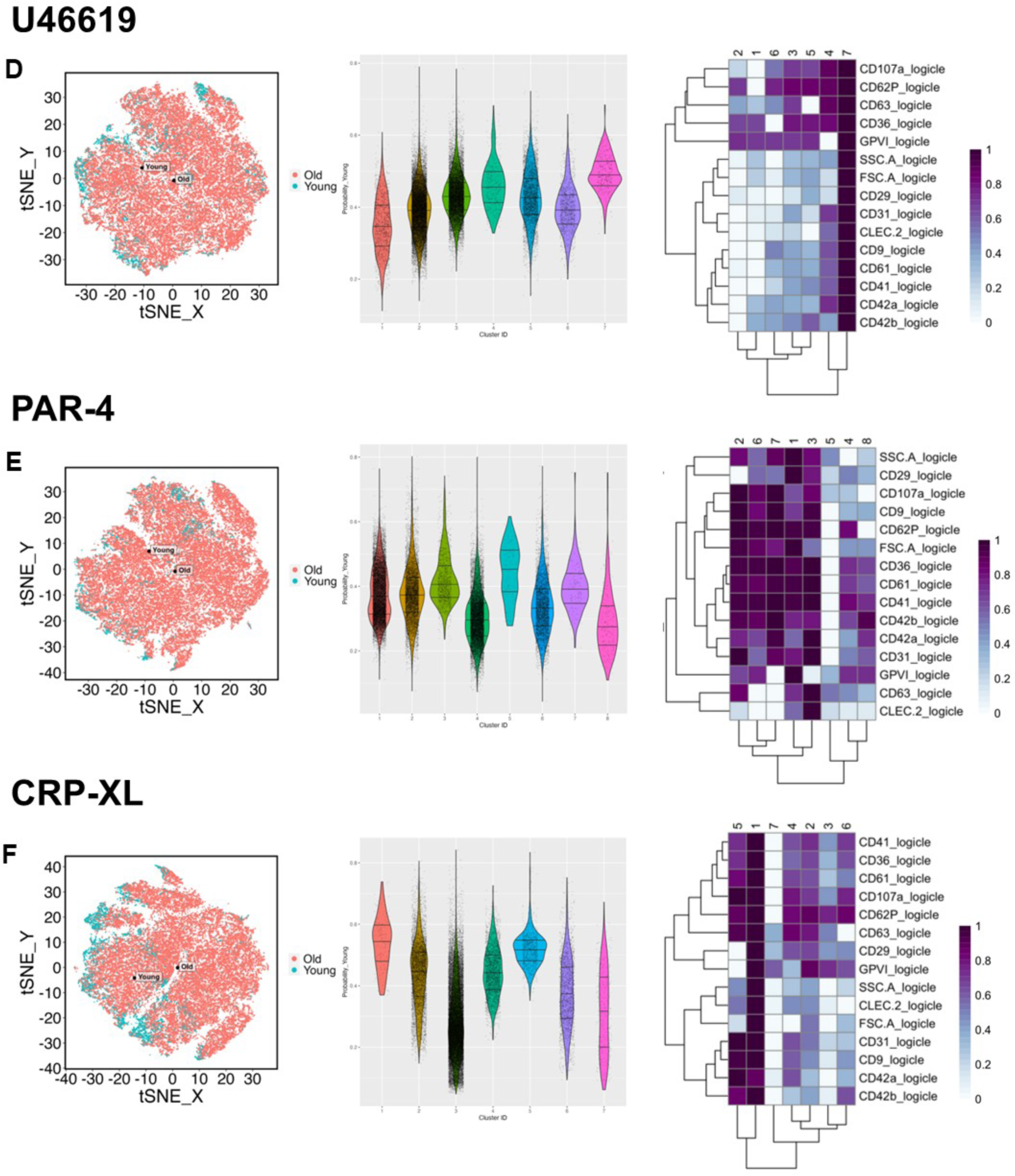
Application of distinguishing markers for young and old platelets weighted by machine learning at above importance 20 to platelet sub-populations using a clustering model (n=20-21 for all) in (A) resting/vehicle-treated, (B) ADP-treated, (C) TRAP-6-treated, (D) U46619-treated, (E) PAR-4-treated, and (F) CRP-XL-treated platelets. Data presented as clustered tSNE population division (left panel), violin plot of density of each cluster (middle panel), and heatmap breakdown of relative marker expression in each cluster (low=white, high=purple; right panel).

## Discussion

The levels of individual platelet surface receptors, basally and following activation, have been well-characterised. However, improvements in flow cytometry and mass cytometry technology now permit much larger antibody panels for simultaneous measurement of receptors on individual platelets. Through greater immunophenotyping of platelets in health and in disease it is increasingly possible to address the question of whether, and what, platelet sub-populations exist. Here we report a 16-parameter phenotypic approach utilising spectral flow cytometry with computational analysis, including machine learning, to phenotype platelets.

Our selection of surface proteins was based on the phenotypic panel used by Blair *et al*.[8] for mass cytometry. First, we set out to measure the effect of different activation pathways of platelet activation on the expression of surface markers. A spectral cytometry-based approach, compared to mass cytometry in terms of antibody costs, machine running costs and processing time, permits an expansion in experimental conditions that can be performed in each run. Across the range of agonists we tested there were significant increases in PAC-1, CD62P, CD63, CD107a, CD61, and CD29 expression, decreases in CD42b expression, and no changes in CLEC-2 expression. These patterns of surface protein changes are consistent with the current understanding of the field and echo those presented by Blair *et al*.[8] and Hindle *et al*.[38]. This initial analysis relies on median fluorescence scores derived from measurements of 10,000 platelets per subject, and comparative variation determined across all 21 donors. Dimensionality reduction (tSNE) and clustering provides for greater analytical weight for each individual platelet across all replicates. It also allows for the visualisation of the contribution of each individual subject to the interpreted expression patterns[39]. This approach validated that these observed patterns are shared across donors, confirming these effects were driven by functional responses and not by potential batch variation.

Unlike previous reports[38], based on P-selectin expression and PAC-1-binding, k-means clustering silhouette analysis within our dataset did not support the presence of 4 (or more) distinct sub-populations. We did observe that across all agonists, on activation the cluster of platelets that had the largest drop in CD42a/b were not the same cluster of platelets that had the highest increase in classic activation markers (CD62P/PAC-1). This implies that platelets are potentially pre-destined to an activation-induced consequence, namely predominantly ‘shedding’ or ‘degranulation’ sub-populations. However, further characterisation of these potential sub-populations would be required to determine their composition pre-activation.

We next turned to machine learning to construct unbiased algorithms to uncover potential classifications at an individual platelet level and to compare predictive capacities between vehicle- and agonist-treated datasets. Consistent with platelet biology and our traditional user-led population analysis, CD62P expression and PAC-1 binding were consistently identified with agonist stimulation, as was loss of CD42b expression. Interestingly, the weighted importance of CD107a, lysosomal-associated membrane protein 1 (LAMP-1), was only high for PAR-1 activation using TRAP-6. Using the computed weightings, machine learning correctly identified over 89% of platelets within the training data set. When applied to an unseen data set, correct identification was predictably lower but maintained greater than 77% accuracy, rising to 91-92% for PAR-stimulated platelets. The difference in predictive capabilities is likely due to U46619 and ADP being comparatively weaker secondary agonists and therefore producing a less uniform pattern of response[40–43].

We next applied the same machine learning approach to differentiate between young and old platelets within mRNA stain-based, flow-sorted platelet samples which were subsequently vehicle- or agonist-stimulated. Strikingly, there is remarkable consistency in the highly-weighted parameters across the unstimulated and stimulated samples with FSC-A, CD41, SSC-A, GPVI, CD61, and CD42b featuring prominently.

A significant change in CD41 with platelet age is consistent with previous work from our group looking at the proteomics and transcriptomics of young and old platelets[33], and from others looking at their thrombotic potential[19]. Similarly, an association between GPVI levels and young platelets have been recently reported by Veninga *et al*.[44] in human platelets and by us using a temporal labelling approach in mice[12].

Interestingly, FSC-A, which is considered an approximate indicator of size in flow cytometry, is also highly important in our machine learning algorithm when distinguishing between young and old platelets. One point of contention concerning the field of platelet ageing is whether platelet size changes with age. Although there is some evidence linking mean platelet volume to thrombotic risk[45,46], there is also contradictory data suggesting that the two variables are independent of one another[12,47,48]. The hypothesis that young platelets are larger was originally proposed in the 1960s[49,50], however by the mid-1980s several studies were published reporting no correlation between platelet age and size[48]. Regardless, a caveat of using SYTO-13 dye as a surrogate marker of platelet age is that larger platelets may have more mRNA, and subsequently take up more dye and appear brighter, skewing sorting and subsequent analysis[51]. However, previous work from our group using a similar mRNA dye noted variations in megakaryocyte-derived mRNAs such that young and old platelets would have to vary 32-64-fold in size for such a relationship to hold.[33] Similarly, it is also notable that SSC-A, generally taken as an indicator of internal complexity (i.e. granularity) of a cell, is a highly important discriminator of young versus old platelets. This finding is directly in line with our observations that platelets lose approximately 50% of their total protein content and mitochondria as they age in the circulation[33]. Similarly, in turn this parameter may also reflect the potential density of a platelet, a measure that studies have suggested is an accurate indicator of platelet age[52–54].

For unseen samples, machine learning-based identification of young and old platelets was 76% accurate in vehicle control samples. This accuracy was maintained in platelets treated with PAR-4 or U46619 but decreased to 67% in platelets treated with ADP. Notably, identification of ADP-stimulated platelets was primarily determined by 4 parameters, whilst for all other conditions 6-7 parameters were used, indicating that perhaps the composition of this panel could be altered for improved accuracy. Conversely, predictive accuracy was higher in samples treated with TRAP-6 or CRP-XL, at 83% and 84%, respectively. In addition, CD62P was more highly-weighted as an additional discriminatory parameter in these samples, which reflects the well-described greater thrombotic potential of young platelets and their higher CD62P degranulation[12,55].

Finally, we applied our machine learning-based identification of young and old platelets back to our healthy platelet populations. This analysis demonstrated that young platelets, as defined by machine learning in each condition, were consistently associated with platelet clusters carrying higher levels of activation markers, in accordance with our earlier reports[12,33]. This suggests that machine learning algorithms employing FSC-A and SSC-A, in addition to surface markers, can be used to further discriminate platelet sub-populations in healthy individuals based on circulatory age.

In conclusion, we present a 16-parameter, flow cytometry-based assay coupled to powerful bioinformatic approaches to undertake unparalleled, deep phenotyping of platelets and their functionality. The incorporation of machine learning into this workflow provides impartial analysis and predictive capability at a by-platelet and by-individual level. We posit that this approach combining surface and physical markers will be highly valuable in phenotyping platelet sub-populations and studying platelet populations in disease or pathological states.

## Acknowledgements

The authors acknowledge the support of the Flow Cytometry Core Facilities at the Blizard Institute. We also thank Prof. Andrew Frelinger and Dr. Benjamin Spurgeon for their valuable advice.

## Authorship contributions

AV and PCA designed the research, performed the assays, collected data, analysed and interpreted data, performed statistical analysis, and wrote the manuscript. JB and HEA designed the research, analysed and interpreted data, performed statistical analysis, and revised the manuscript. JB performed bioinformatic analyses. CAM and TDW designed the research, analysed and interpreted data, and revised the manuscript.

## Supplemental Data

**Supplemental figure 1:**
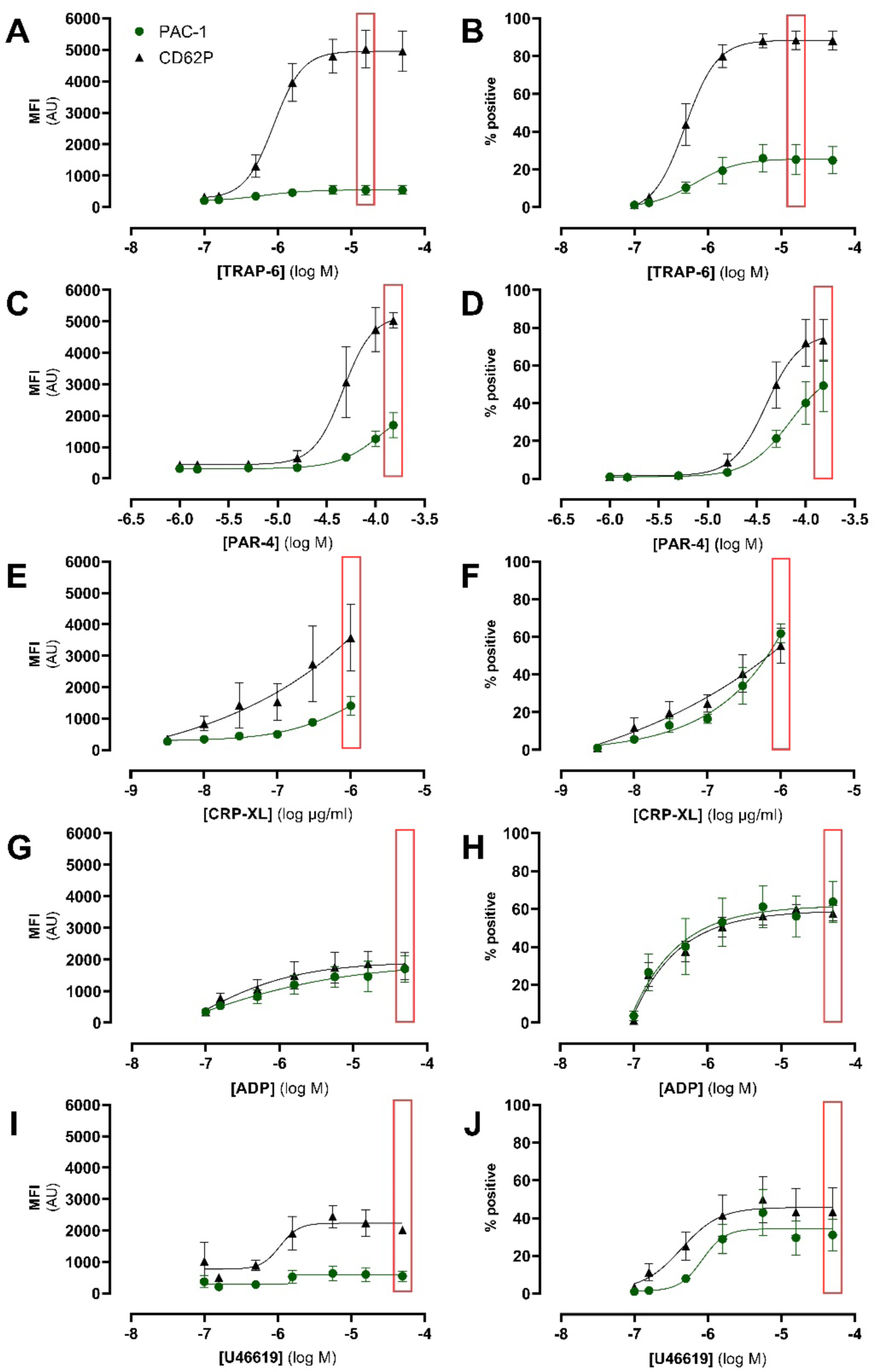
Concentration-response curves of platelet activation in response to (A) TRAP-6 (0.1-10 µM), (B) PAR-4 (3-100 µM), (C) CRP-XL (0.03-3 µg/ml), (D) ADP (0.3-30 µM), and (E) U46619 (0.1-10 µM) measured by PAC-1 binding and CD62P exposure. Agonist concentrations chosen for subsequent assays highlighted within red boxes. Data are shown as MFI ±SEM or % positive ±SEM (n=3-4) and analysed using non-linear regression.

**Supplemental figure 2:**
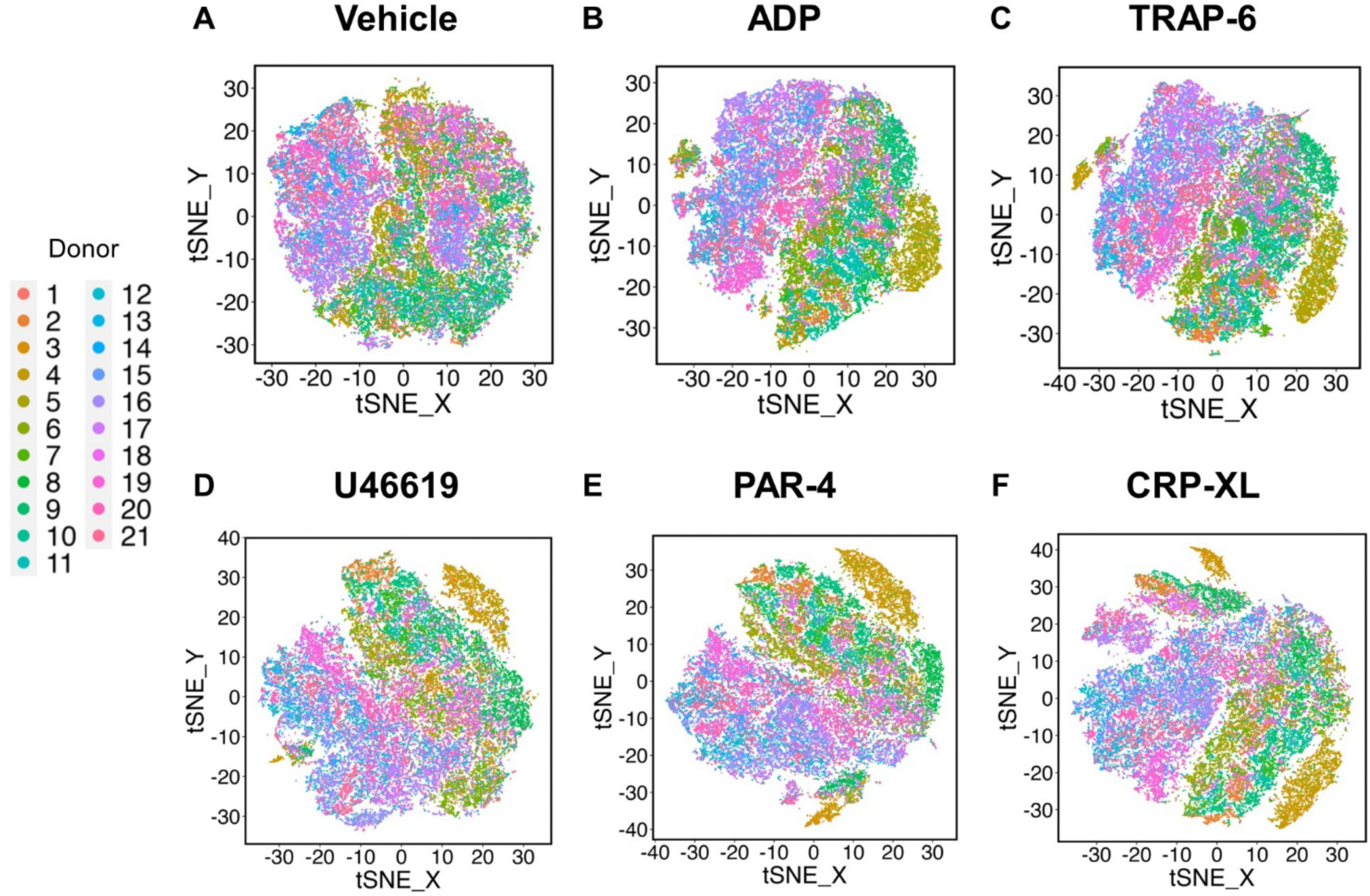
Composite tSNE of all donor platelets (n=20-21) for (A) vehicle, (B) ADP, (C) TRAP-6, (D) U46619, (E) PAR-4, and (F) CRP-XL.

**Supplemental figure 3:**
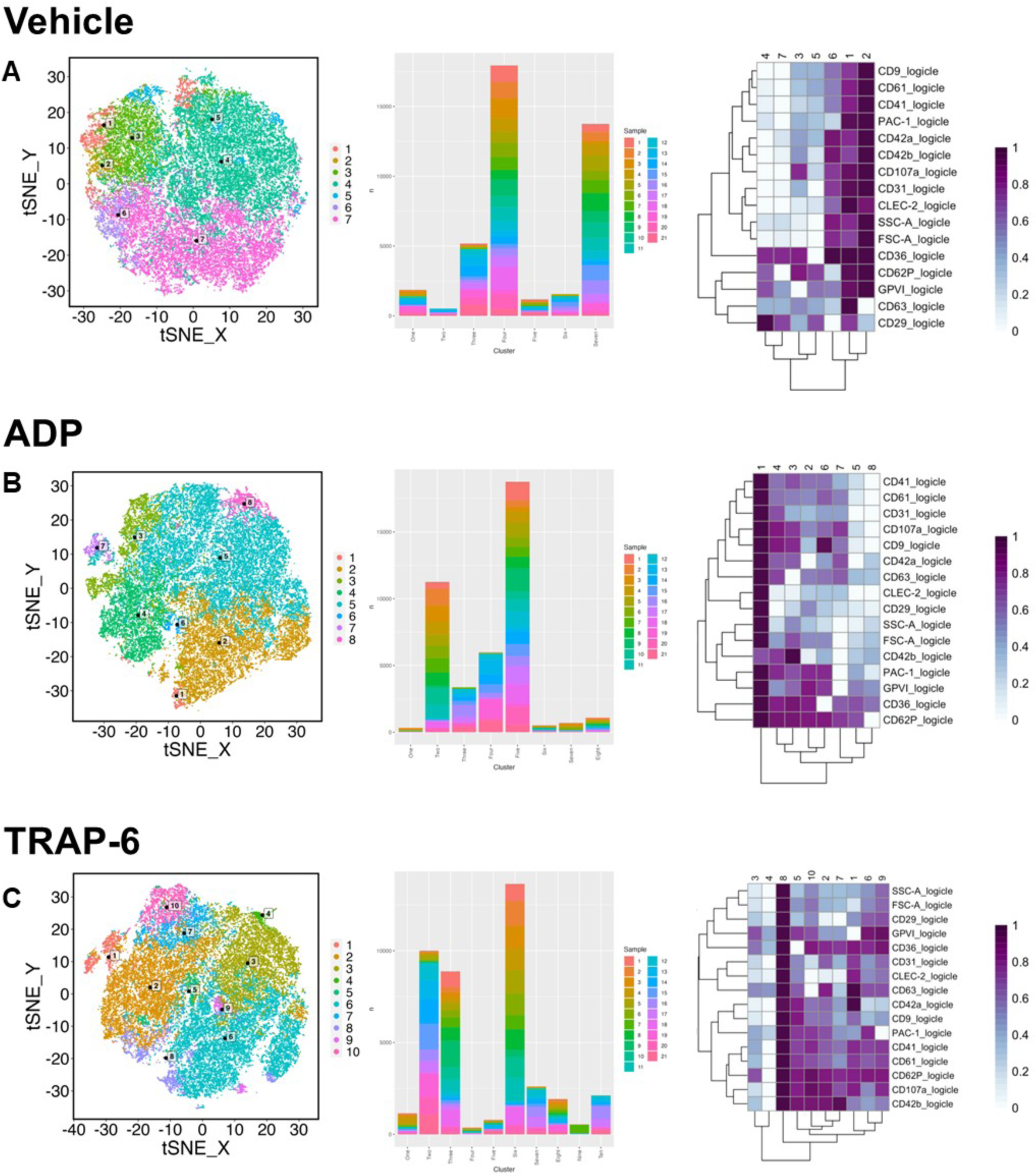
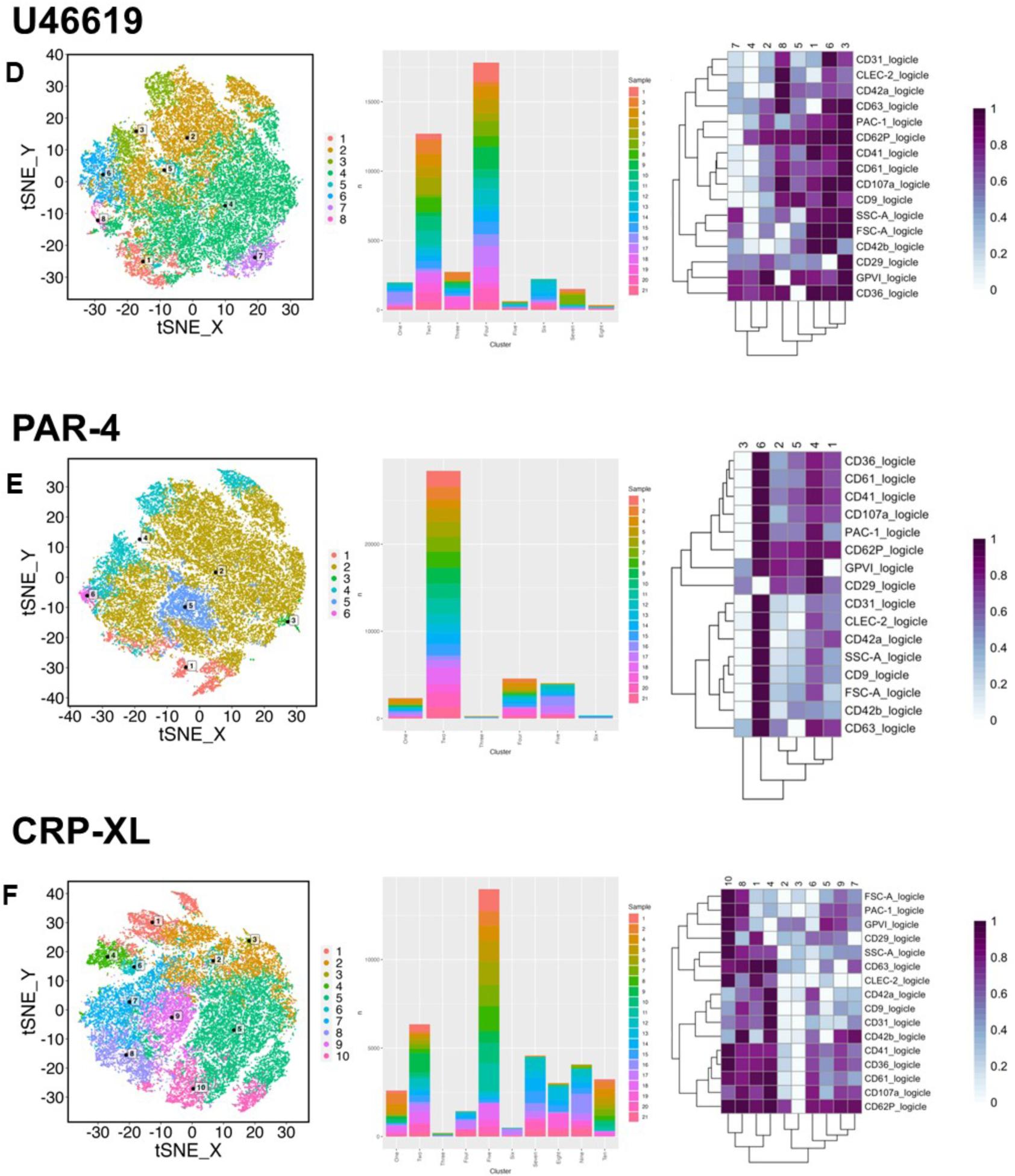
High dimensionality analysis of platelet sub-populations using a clustering model in (A) resting/vehicle-treated, (B) ADP-treated, (C) TRAP-6-treated, (D) U46619-treated, (E) PAR-4-treated, and (F) CRP-XL-treated platelets. Data presented as clustered tSNE population division (left panel), corresponding donors making up each cluster (middle panel), and heatmap breakdown of fold-change in marker expression in each cluster (low=white, high=purple; right; right panel; n=20-21).

**Supplemental figure 4:**
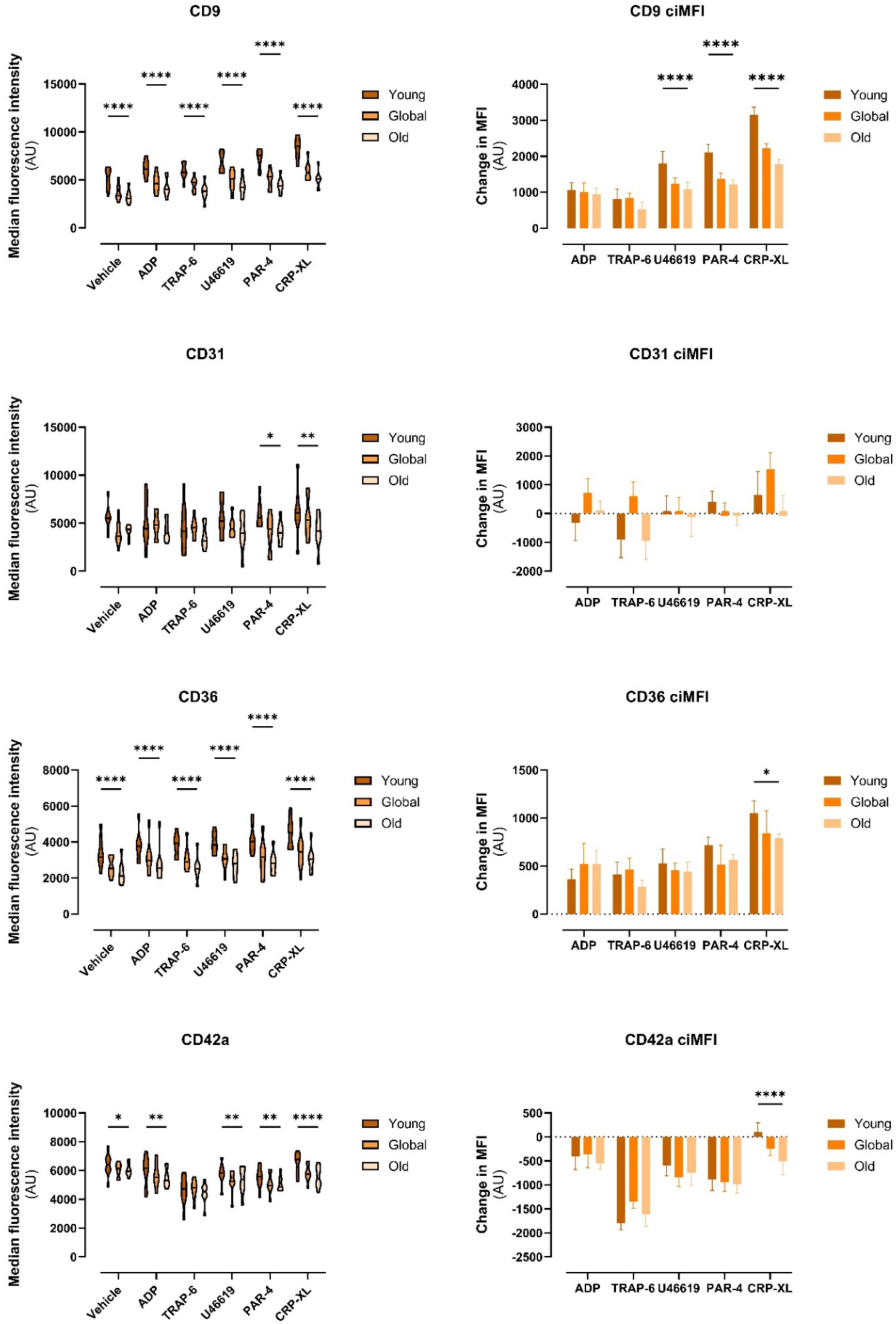
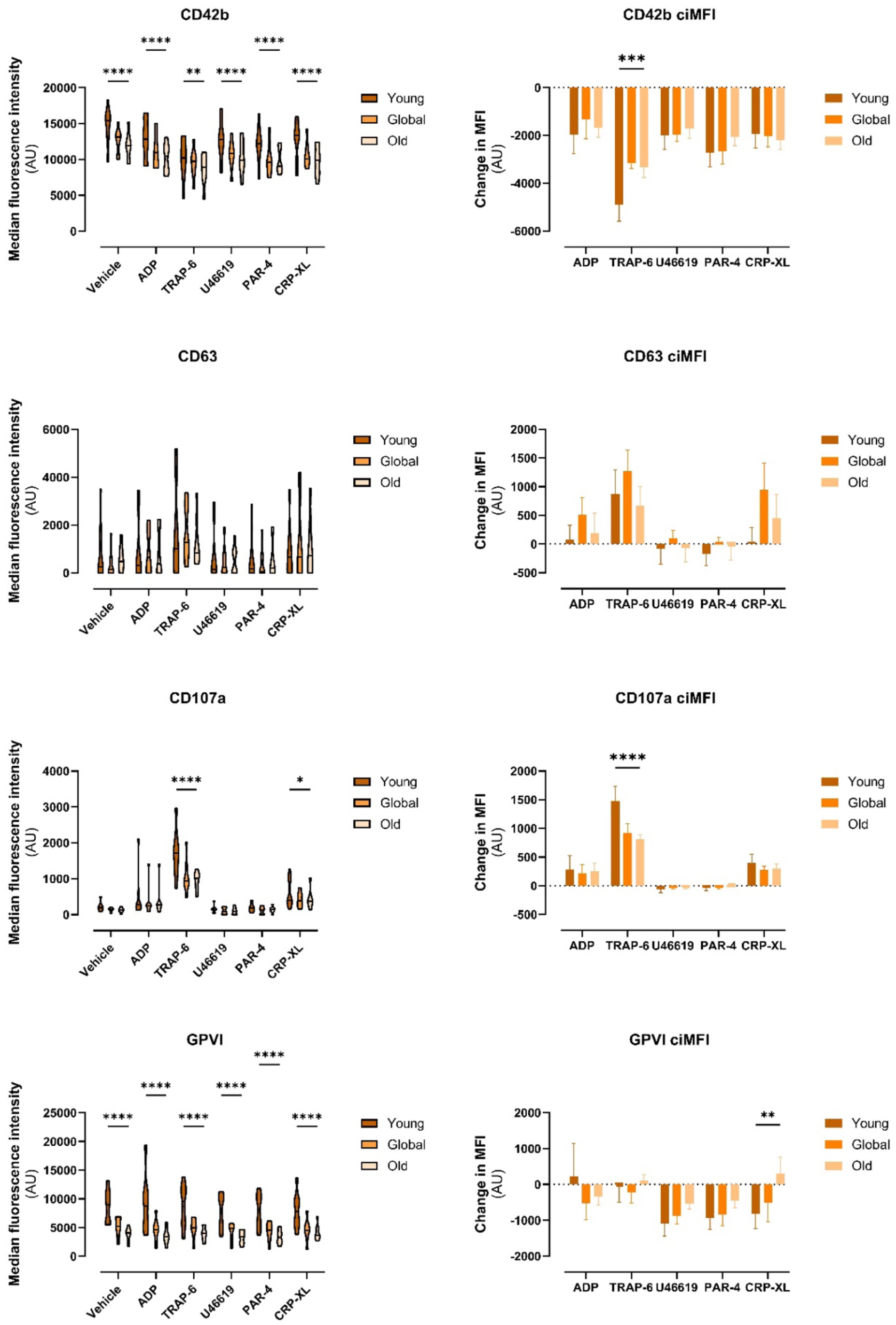
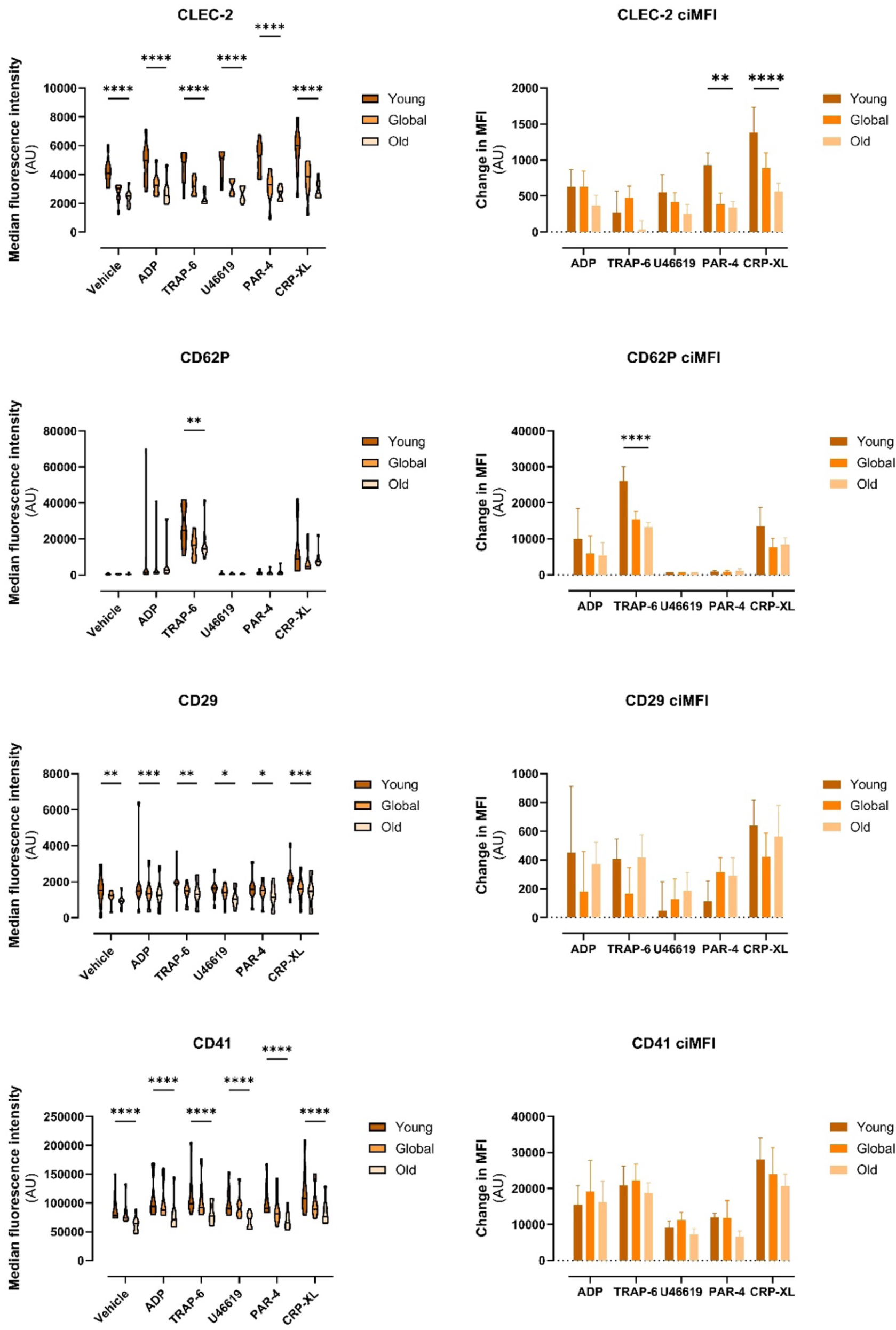
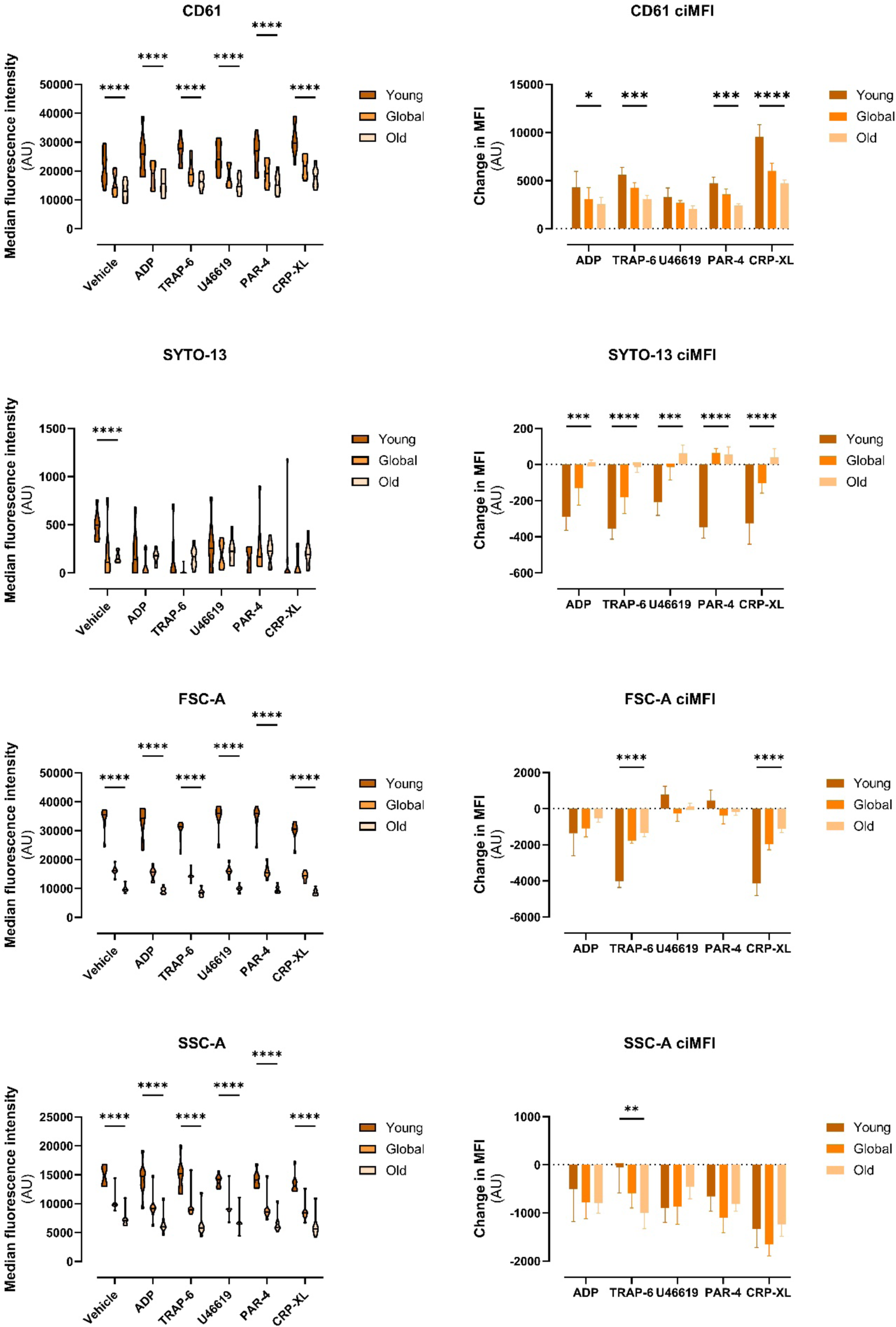
Changes in surface marker expression in “young” and “old” resting and activated platelets expressed as raw MFI (left) and change in MFI (right). Data are shown as MFI ±SEM (n=8) and analysed using 2-way ANOVA or mixed-effects analysis with Tukey’s multiple comparisons test.

**Supplemental table 1:** Details of panel antibodies.

| Marker | Alias | Role | Fluorophore | Clone | Dilution factor | Laser (Channel) | Supplier (Catalogue number) |
| --- | --- | --- | --- | --- | --- | --- | --- |
| CD107a | LAMP-1 | Lysosome degranulation | PE | H4A3 | 1:200 | Yellow/Green (YG1) | BioLegend (328608) |
| CD29 | GPIIa | Adhesion integrin | BUV737 | MAR4 | 1:200 | Ultraviolet (UV14) | BD Biosciences (748618) |
| CD31 | PECAM-1 | α-granule protein; platelet-endothelial cell interaction | BV785 | WM59 | 1:200 | Violet (V15) | BioLegend (303148) |
| CD36 | GPIV | α-granule protein; Scavenger receptor | BV605 | CB38/NL 07 | 1:70 | Violet (V10) | BD Biosciences (563518) |
| CD41 | GPIIb, | Fibrinogen binding integrin | APC-Cy7 | HIP8 | 1:100 | Red (R7) | BioLegend (303716) |
| CD61 | GPIIIa, |  | BUV395 | RUU-PL7F12 | 1:70 | Ultraviolet (UV2) | BD Biosciences (745717) |
| CD42a | GPIX | Adhesion integrin; binds vWF | eFluor 450 | GR-P | 1:30 | Violet (V3) | Thermo Fisher Scientific (48-0428-42) |
| CD42b | GPIbα | | BV650 | HIP1 | 1:30 | Violet (V11) | BioLegend (303926) |
| CD62P | P-selectin | α-granule protein; platelet-endothelial/ leukocyte binding | BB700 | AK-4 | 1:100 | Blue (B9) | BD Biosciences (566565) |
| CD63 | LAMP-3 | δ-granules, lysosomes marker | PE-Cy7 | H5C6 | 1:70 | Yellow/Green (YG9) | Thermo Fisher Scientific (25-0639-42) |
| CD9 | TSPAN29 | Tetraspanin; modulate collagen response/ GPIIb/IIIa | BV510 | M-L13 | 1:70 | Violet (V7) | BD Biosciences (563640) |
| CLEC-2 | CLEC-1b | Thromboinflammatory receptor | BV711 | 219133 | 1:70 | Violet (V13) | BD Biosciences (748142) |
| GPVI |  | Collagen receptor | AF647 | HY101 | 1:200 | Red (R2) | BD Biosciences (564701) |
| PAC-1 |  | Binds activated confirmation of receptor GPIIb/IIIa integrin | FITC | PAC-1 | 1:10 | Blue (B2) | BD Biosciences (340507) |

